# ProtBLIP2-SST: Protein Function Prediction via BLIP2 with Sequence, Structure, and Text

**DOI:** 10.64898/2026.07.10.737551

**Authors:** Zhuoyang Chen, Qiong Luo

## Abstract

Protein function prediction traditionally relies on structured gene ontology (GO) labels or multi-label classifiers. However, these labels or classifiers cannot flexibly describe molecular function, biological process, cellular component, and free-text functional narratives in a single output. In comparison, generation-based approaches offer an intuitive paradigm for flexible free-text protein annotation, with large language models (LLMs) as a representative method for protein-text modeling. Recent efforts on utilizing LLMs for protein semantic understanding and annotation generation have adopted sequence-only encoding or sequence-text contrastive alignment paradigms, yet without explicit consideration of three-dimensional structural information. To address these limitations in current protein function prediction methods, we present ProtBLIP2-SST, a two-stage framework built on the BLIP2 model architecture that bridges protein sequence, structure, and text for open-ended protein functional caption generation. Specifically, we first integrate sequence and structure information through SaProt, a protein language model (PLM) with a structure-aware vocabulary that fuses residue tokens with Foldseek-derived 3Di structural tokens. To empower the LLM to understand protein semantics, we employ a Q-Former (a querying transformer in BLIP2) with learnable query tokens as the cross-modal projector to align protein features from the frozen SaProt encoder and text features from a frozen BiomedBERT via protein–text contrasting, protein–text matching, and protein captioning objectives. After alignment, the protein features are linearly projected and prepended to the prompt embeddings of the LLM for protein captioning fine-tuning with LoRA. Trained on 441k protein–text pairs from Swiss-Prot with corresponding structures from the AlphaFold Database, our ProtBLIP2-SST outperforms sequence-only and sequence-text alignment baselines on protein captioning metrics, with ablation studies demonstrating the effectiveness of integrating structure with sequence information for improved protein understanding. Through a unified two-stage alignment-and-generation pipeline, ProtBLIP2-SST integrates protein sequence and structural information, overcomes the rigidity of traditional GO-centric classification, generating open-ended captions that jointly describe molecular function, subcellular location, and homology context in one single output.

## I. Introduction

Computational protein function prediction is dominated by GO-centric multi-label classification [1, 2]. Although many deep learning methods that concatenate embeddings from protein language models (PLMs) with various prediction heads [3, 4] have been proposed for accurate prediction on curated functional labels, they cannot produce novel textual hypotheses, combine function with homology or subcellular context in one response, or leverage the rich free-text annotations in UniProt/Swiss-Prot [5] (Figure 1). Large language models (LLMs) excel at biomedical text understanding [6] but cannot directly interpret protein data: amino acid sequences, despite being composed of single-letter symbols in the alphabet, encode biochemical properties rather than linguistic semantics, and three-dimensional structural conformations or molecular interaction patterns cannot be fully converted to natural language descriptions. Recent efforts on utilizing LLMs for protein semantic understanding and annotation generation have adopted sequence-only encoding [7] or sequence-text contrastive alignment paradigms [8, 9, 10, 11], yet without explicit consideration of three-dimensional structural information. Structure-aware [12, 13] or multimodal PLMs [14, 15, 16] have been derived but have not been utilized by LLM for text generation.

**Fig. 1.**
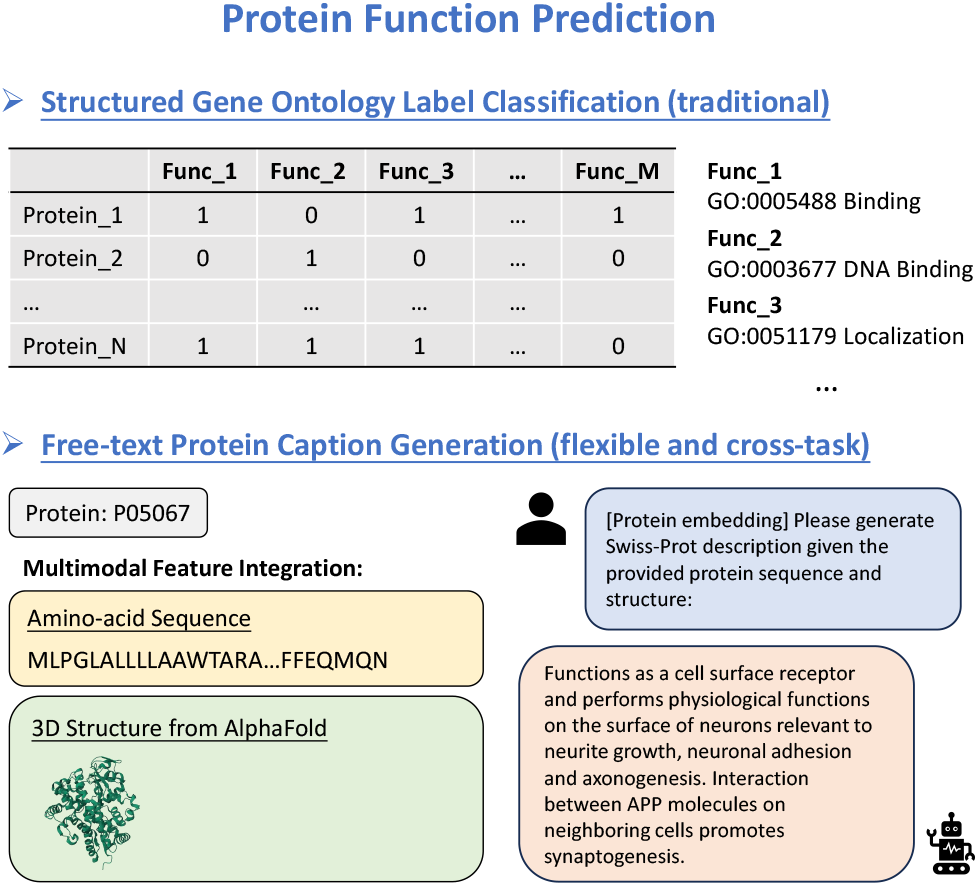
Examples of free-text protein caption generation compared to the traditional structured GO label supervised classification task.

To unify sequence, structure, and text under one generative framework and treat protein function prediction as protein-to-text captioning, we propose ProtBLIP2-SST: A protein function annotation generator with **S**equence, **S**tructure, and **T**ext in the BLIP-2 framework [20]. Captions follow Swiss-Prot field structure (FUNCTION, SUBCELLULAR LOCATION, and SIMILARITY), framing function prediction as conditional text generation rather than fixed-label classification, which makes functional semantics explicit and reusable across downstream tasks. Our model adopts a two-stage training process to enhance protein-function relationship modeling. The first stage is a contrastive learning framework the same as in BLIP-2. We first encode protein sequence and structure retrieved from the Alphafold database [21] together with SaProt [14], a model with structure-aware vocabulary that integrates residue tokens with structure tokens [22]. Then we train a cross-modal projector, Q-Former, to align the protein embeddings with the space of text embedding from the target LLM through contrastive, matching, and captioning objectives. This stage serves both to bridge the modality gap and to warm up the representation before LLM fine-tuning. The second stage is fine-tuning the scientific LLM Galatica-1.3B [7] on the Swiss-Prot dataset. The protein embeddings output by the Q-Former are linearly projected and prepended to the prompt embeddings of the LLM, and the model is trained for protein captioning via next-token prediction.

Our results show that ProtBLIP2-SST outperforms recent advanced models on both protein–text retrieval and protein captioning tasks, demonstrating the effectiveness of the introduction of structural information and the integration with sequences. Further analysis reveals that structural information benefits FUNCTION descriptions most, while offering marginal gains on SIMILARITY family labels.

The contributions of this work are summarized as follows:

- We introduce ProtBLIP2-SST, a unified framework for open-ended protein function annotation that integrates sequence, structure, and text modalities through a BLIP2-style two-stage alignment and generation pipeline.
- We establish systematic benchmarks for stage1 retrieval and stage2 captioning on the Swiss-Prot test set, demonstrating that ProtBLIP2-SST (650M) achieves the strongest performance among variants trained under identical procedures.
- We perform field-stratified analysis showing that structural information improves FUNCTION prediction most significantly, but offers limited gains for SIMILARITY-based annotations, clarifying the task-dependent value of 3D structure in protein captioning.

## II. Related Work

### A. Protein language models

Sequence-only PLMs such as Esm2 [23] are trained with masked language modeling (MLM) on amino-acid sequences. Researchers have also explored incorporating structural information as an additional constraint. ISM is first pre-trained with the MLM objective and subsequently fine-tuned to predict structure tokens [13]. ProstT5 treats sequence and structure as a bilingual T5 translation task, converting amino-acid tokens and structure tokens bidirectionally [12]. SaProt augments the vocabulary with structure-aware tokens derived from Foldseek’s 3Di alphabet [14]. ProTrek performs tri-modal contrastive learning for protein-text-structure retrieval without open-ended generation [16]. ESMC adopts a three-track architecture for sequence, structure, and function, pre-trained with the MLM objective [15]. As discussed above, models that concatenate PLM embeddings with prediction heads for structured-label classification lack the flexibility to express free-form functional descriptions and novel biological functions.

### B. Vision-language pre-training

Early vision-language pre-training methods use dual-encoder architectures such as CLIP [24], which employ cross-modal contrastive learning to align image and text representations in a shared embedding space. These models are good at retrieval tasks but perform poorly on text generation, as their contrastive objective does not train the model to condition text generation on visual inputs. Subsequent work try to unify vision-language understanding and generation. BLIP [25] introduces a bootstrapping framework with an image-grounded text encoder and decoder, jointly optimized via image-text contrastive learning, image-text matching, and image-grounded text generation. However, BLIP requires end-to-end training of large-scale multi-modal transformers, leading to substantial computational costs and making it difficult to utilize novel advanced unimodal pre-trained models. BLIP-2 [20] combines frozen image encoders and LLMs with a lightweight Q-Former trained for representation learning then generative conditioning. We replace the image encoder with a frozen PLM and the LLM with Galactica in ProtBLIP2-SST and then fine-tune it with mid-layer Low-Rank Adaptation (LoRA) [26].

### C. Protein–text modeling

Several studies adapt the vision-language pre-training frameworks for protein–text modeling. ProtST [8] and ProtCLIP [9] align protein sequences with text for retrieval using CLIP. ProtT3 utilizes Q-Former for sequence-text alignment and Galactica fine-tuning for captioning and Question-Answering tasks [11]. Prot2Text-V2 uses H-SCALE and a large instruction-tuned decoder for enhanced alignment and text generation, respectively [10]. ProteinChat encodes protein structure and directly uses its embeddings without alignment with text for downstream LLM fine-tuning [17]. More recent work has explored multi-level structural integration. ProtChat-GPT [19] conducts separate encoding for each structural level and aligns them sequentially: first sequence-to-text via its PLP-Former, then structure-to-sequence via a gating module, which potentially limits effective interaction between sequence and structural features. In contrast, Prot2Chat [18] jointly encodes sequence and structure through ProteinMPNN [27] and then fuses question embeddings with protein features by a text-aware adapter. While Prot2Chat achieves efficient early fusion, its graph-based encoder is designed for structure-conditioned sequence generation rather than general-purpose protein representation, and its text-aware module lacks explicit contrastive alignment between protein and text embedding spaces.

Compared to existing work, ProtBLIP2-SST addresses these limitations through a unified structure-aware representation and explicit multi-objective alignment (Table I). We use SaProt as the protein encoder, which fuses sequence and structural information into a single vocabulary of structure-aware tokens, avoiding separate encoders or sequential alignment stages. This design ensures that sequence and structural features interact from the earliest representation layer, overcoming the limited cross-modal interaction in ProtChatGPT’s progressive pipeline and the non-generalizable encoding of Prot2Chat’s ProteinMPNN module.

**TABLE I.** Comparison of protein–text modeling methods. ✓: evaluated; ×: not evaluated; ?: incomplete. Modality: S = sequence; St = structure; T = text; S+St (Sep) = separate encoders; S+St (Uni) = unified Sa-vocab (SaProt). Integ. = integration (Lin: Linear; mlp; qf: Q-Former). Align. = alignment (CL: contrastive; Gen: Generative; PT: pretraining). Tasks: Gen. = captioning/QA; Ret. = retrieval.

| Method | Modality | Integ. | Align. | Gen. | Ret. |
| --- | --- | --- | --- | --- | --- |
| *Contrastive-only* |  |  |  |  |  |
| ProtST [8] | S | Lin | CL | × | ? |
| ProtCLIP [9] | S | Lin | CL | × | ? |
| *Generative-only / single-stage* |  |  |  |  |  |
| Galactica [7] | T | — | PT | ✓ | × |
| ProteinChat [17] | St | Lin | Gen | ? | × |
| Prot2Chat [18] | S+St (Sep) | MLP | Gen | ✓ | × |
| ProtChatGPT [19] | S+St (Sep) | MLP | Gen | ✓ | ✓ |
| *Two-stage alignment & generation* |  |  |  |  |  |
| Prot2Text-V2 [10] | S | MLP | CL+Gen | ✓ | × |
| ProtT3 [11] | S | QF | CL+Gen | ✓ | ✓ |
| **Ours, ProtBLIP2-SST** | **S+St (Uni)** | **QF** | **CL+Gen** | **✓** | **✓** |

## III. Methods

### A. Model Architecture

ProtBLIP2-SST adopts a two-stage training paradigm that bridges the modality gap between protein sequences/structures and natural language descriptions (Figure 2). The architecture comprises three principal components: a frozen protein language model (PLM) for sequence encoding, a lightweight cross-modal projector for protein–text alignment, and a frozen large language model (LLM) connected with parameter-efficient fine-tuning adapters for conditional text generation.

**Fig. 2.**
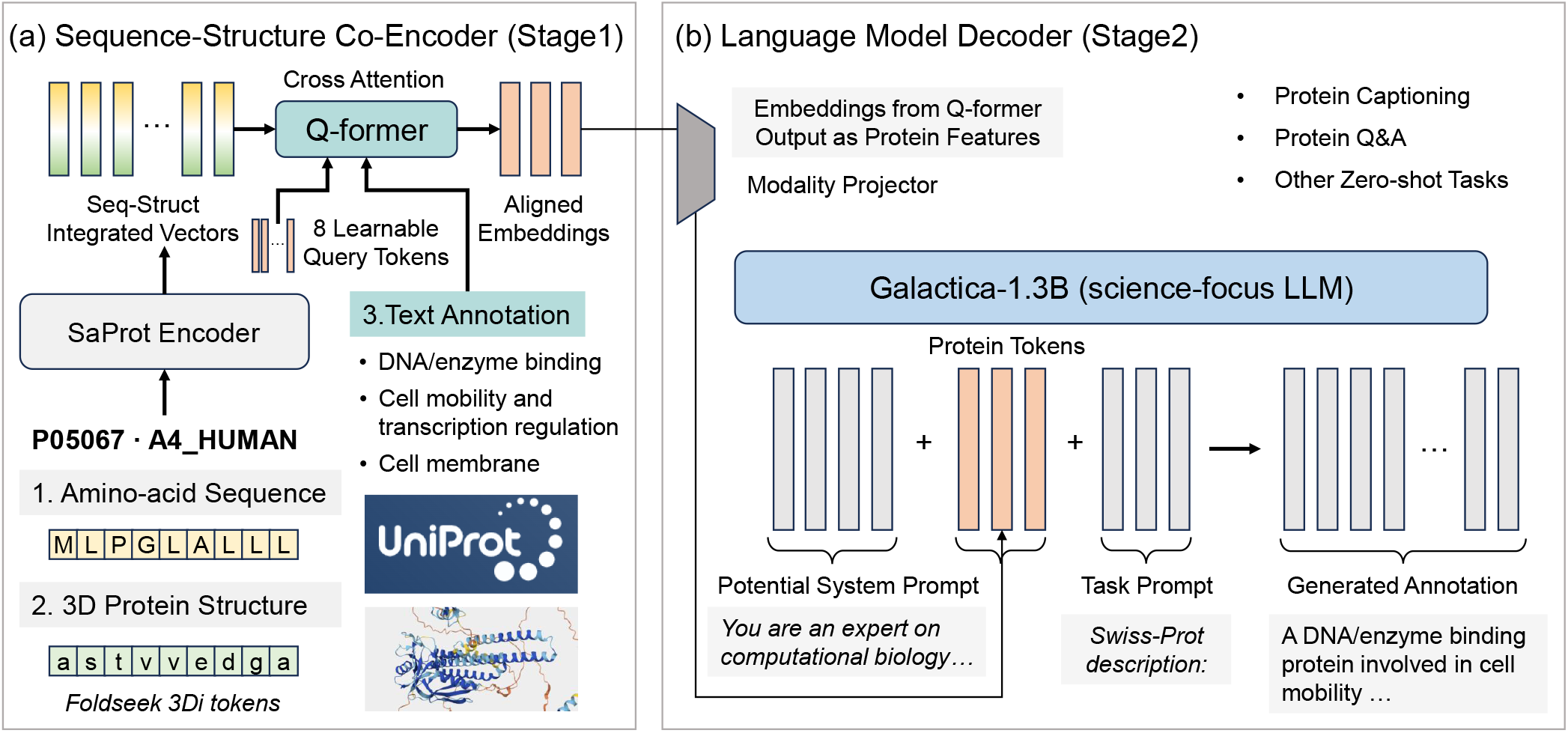
Schematic illustration of the ProtBLIP2-SST architecture. **(a):** In the protein sequence-structure co-encoder, the amino-acid sequence and its 3D structure are first processed and co-encoded by SaProt. The 8 learnable query tokens and the text annotation are passed through the same text encoder in Q-former. Then cross-attention is performed between the 8 query tokens and sequence-structure integrated embeddings (as keys and values). The output embeddings with the same length of 8 with query tokens are viewed as protein features and further used for contrastive learning (alignment). **(b):** The aligned embeddings from the stage1 training are further projected to share the same dimension with Galactica-1.3B and prepended to the task prompt embeddings for protein captioning fine-tuning.

1. *Protein Encoding:* Given an input protein sequence **s** = [*s*_1_, *s*_2_, …, *s*_*L*_] of length *L* ≤1024, a frozen PLM *f*_*θ*_ computes contextualized residue embeddings:

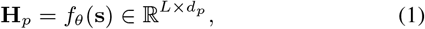

where *d*_*p*_ denotes the PLM hidden dimension (*d*_*p*_ = 480 for 35M-parameter encoders; *d*_*p*_ = 1280 for 650M-parameter encoders; *d*_*p*_ = 1024 for ProstT5 encoder). We use three categories of protein encoders to investigate the contribution of structural information: (1) Esm2 (sequence-only) 35M and 650M, which is adopted in ProtT3. (2) SaProt (*SaProt_35M_AF2* and *SaProt_650M_AF2*) that used in our ProtBLIP2-SST model. (3) ProstT5 (sequence-structure bidirectional translation) with 1.5B parameters. All PLM parameters remain frozen throughout both training stages to preserve pretrained biochemical knowledge and mitigate catastrophic forgetting.
2. *Cross-Modal Projector (Q-Former):* Following the design principle of BLIP-2 [20], we employ Q-Former as the cross-modal projector. We set Q-Former to have *N*_*q*_ = 8 learnable query tokens 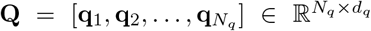, initialized from weights of the BiomedBERT-Abstract text encoder [28] (Figure 3). The protein transformer sub-module of the Q-Former processes these queries through alternating self-attention and cross-attention layers, where cross-attention is inserted every other transformer block to enable interaction with the frozen PLM outputs **H**_*p*_. The output protein queries are denoted as:

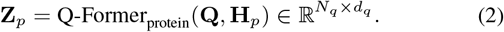

Concurrently, a text transformer sub-module processes input text annotations with a prepended [CLS] token, producing text representations 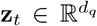 from the [CLS] output. Both submodules share self-attention parameters to facilitate protein– text interaction, with objective-specific attention masking strategies controlling the degree of cross-modal information flow.
3. *Language Model Decoder*.: For conditional text generation, we employ Galactica-1.3B [7], a decoder-only transformer pretrained on large-scale scientific corpora. A linear projection layer 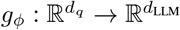 maps the Q-Former outputs to the embedding space of Galactica:

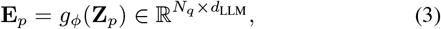

where *d*_LLM_ = 2048. The projected protein embeddings **E**_*p*_ are prepended to the prompt embeddings to condition the LLM on protein-specific information.

**Fig. 3.**
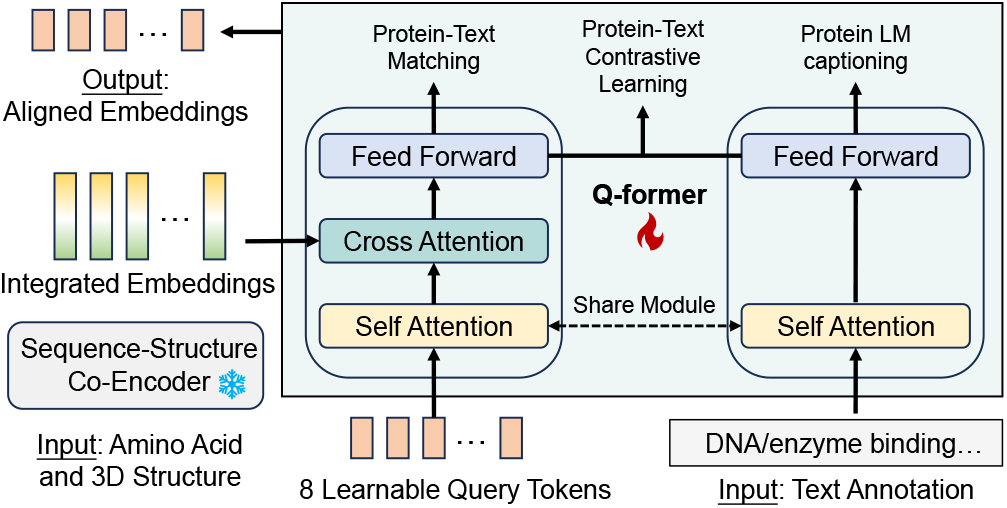
Overview of the Q-former architecture. Q-former takes the text annotation, amino-acid sequence, and 3D structure of a protein as inputs. Eight learnable query tokens are setup to capture protein features by cross attention with integrated embeddings from the sequence-structure co-encoder SaProt. BiomedBERT is used as text encoder for both query tokens and text annotations. Three objectives, protein-text contrastive learning, protein-text matching, and protein LM captioning, are calculated between protein features and text features. The aligned embeddings output from the protein part are used as protein features for further LLM fine-tuning.

To enable efficient adaptation while preserving the LLM’s pretrained linguistic capabilities, we apply LoRA to selected weight matrices. We set *r* = 8, *α* = 16, and dropout = 0.1, applying LoRA to query, key, value, and output projections of self-attention modules (*q_proj, k_proj, v_proj, out_proj*) as well as feed-forward layers (*fc1, fc2*). Base LLM weights remain frozen during training.

### B. Stage 1: Protein–Text Representation Learning

The first training stage empowers the Q-Former to extract protein features that are most informative for textual descriptions, following the multi-objective pretraining strategy of BLIP-2. We jointly optimize three complementary objectives: protein–text contrastive learning (PTC), protein–text matching (PTM), and protein captioning (PCap).

1. *Protein–Text Contrastive Learning (PTC):* PTC enforces alignment between protein and text representations via cross-modal contrastive learning. Given a batch of *N* protein– text pairs 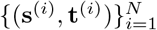, we compute protein representations 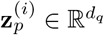by mean-pooling the Q-Former protein queries, and text representations 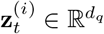 from the BERT [CLS] output. The symmetric InfoNCE loss with temperature *τ* = 0.1 is defined as:

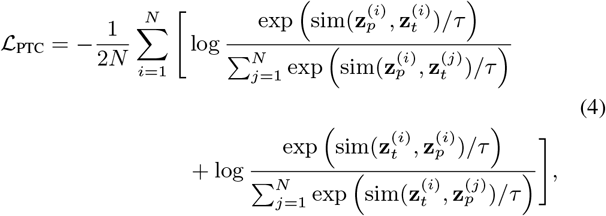

where 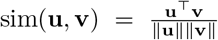 denotes cosine similarity. A unimodal self-attention mask prevents direct interaction between protein queries and text tokens, compelling the queries to extract discriminative protein features from the PLM [11].
2. *Protein–Text Matching (PTM):* PTM formulates cross-modal alignment as a binary classification task to capture fine-grained token-level interactions. A bidirectional self-attention mask allows protein queries and text tokens to attend to each other throughout all layers. The output query embeddings are mean-pooled and fed into a linear classifier 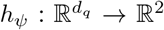to predict whether a pair is matched or unmatched:

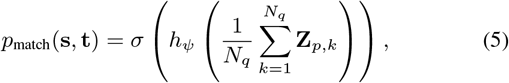

where *σ*(·) denotes the sigmoid function. For each positive pair (**s**^(*i*)^, **t**^(*i*)^), we sample *M* hard negative pairs by replacing either the protein or text with randomly selected alternatives 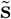 or 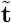 from the batch, respectively. The PTM loss is:

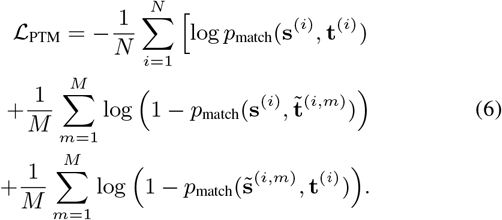
3. *Protein Captioning (PCap):* PCap trains the Q-Former to generate textual descriptions conditioned on protein inputs. A multimodal causal self-attention mask is applied: protein queries attend bidirectionally to each other but not to text tokens; each text token attends to all protein queries and preceding text tokens. This design ensures that text generation relies exclusively on protein features extracted by the queries, functioning as an information bottleneck [11, 20]. Let **t** = [*t*_1_, *t*_2_, …, *t*_*T*_ ] denote a caption of length *T* . The autoregressive language modeling objective maximizes the conditional log-likelihood:

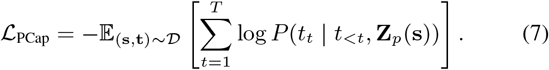

The stage1 training objective combines the three losses with equal weighting:

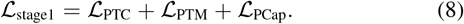
4. *Retrieval Inference:* For protein–text retrieval, we employ a two-stage cascade: (1) PTC embeddings are used for coarse candidate retrieval via maximum inner product search; (2) PTM scores re-rank the top-*K* candidates (*K* = 128) for refined alignment. We report accuracy (Acc) and recall at rank 20 (Rec@20) for both protein-to-text (P2T) and text-to-protein (T2P) directions, evaluated on both in-batch samples and the held-out test set.

### C. Stage 2: Protein–Text Conditional Generation

The second training stage connects the pretrained Q-Former to the frozen Galactica-1.3B decoder to enable protein-conditioned caption generation. The Q-Former weights are initialized from the stage1 checkpoint, providing a warm-start for cross-modal conditioning [11].

Given a protein sequence **s**, the model constructs an input sequence by prepending the projected protein queries **E**_*p*_ to the prompt tokens **p** = [*Swiss-Prot description:*] and caption tokens **t**:

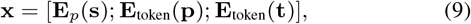

where **E**_token_(·) denotes the token embedding lookup. The language modeling loss is computed exclusively over caption tokens:

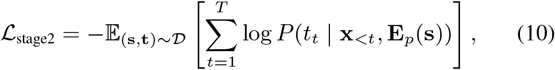

with gradients propagated through the LoRA adapters and Q-Former parameters while the PLM and LLM base weights remain frozen.

During inference, we employ beam search with beam width 5 and maximum generation length of 128 new tokens. The prompt *Swiss-Prot description:* is prepended to condition the model on the captioning task.

## IV. Experiments

### A. Datasets

We supplement the Swiss-Prot V3 dataset that was used in ProtT3 [11] with predicted 3D structures from the AlphaFold Database v4 [21]. A protein record is retained only when its AlphaFold prediction is available, yielding paired sequence– structure–text triplets. Each record comprises the amino-acid sequence, the 3D structure, and a composite functional description concatenating *FUNCTION, SUBCELLULAR LOCATION*, and *SIMILARITY* annotation fields from Swiss-Prot. We then filter records that miss any aspect of the triplet. The final filtered dataset comprises 410,925 training, 9,236 validation, and 9,239 test proteins. During training, protein sequences are truncated to *L* ≤1024 residues and captions to *T*≤ 128 tokens.

### B. Performance Metrics

1. *Stage 1: Protein–Text Retrieval:* We evaluate retrieval performance using accuracy (Acc) and Recall@20 after PTM re-ranking over 128 candidates. Metrics are averaged across protein-to-text and text-to-protein directions for both in-batch retrieval (batch size 64) and test-set retrieval (full 9,239 samples).
2. *Stage 2: Caption Generation:* We assess generation quality using standard natural language generation metrics computed against ground truth captions on the Swiss-Prot test set as in ProtT3 [11]. Exact Match (EM) measures the proportion of predictions identical to references; BLEU-2/4 is the n-gram precision with brevity penalty; ROUGE-1/2/L is the 1/2-gram recall and longest common subsequence, respectively; METEOR is the harmonic mean of unigram precision and recall with synonym matching.

### C. Environment Setup and Parameters

All experiments are conducted on a server equipped with two Intel Xeon Platinum 8375C CPUs (64 cores / 128 threads aggregate), 503 GiB system memory, and two NVIDIA A100 80GB PCIe GPUs. Training employs the AdamW optimizer with initial learning rate 10^−4^, weight decay 0.05, and a linear warmup cosine decay schedule (1,000 warmup steps, minimum learning rate 10^−5^) for both stages. *Stage1:* The batch size is 32 for PTC/PCap and 64 for PTM (hard negative mining). Training proceeds for 50 epochs with seed 42. The Q-Former employs *N*_*q*_ = 8 query tokens. *Stage2:* The batch size is 32 for training and 4 for inference. Training spans 10 epochs with LoRA. Maximum sequence lengths are 128 tokens for captions and 1024 residues for proteins. The checkpoints of epoch 10 are saved and evaluated for models in stage2.

### D. Baseline methods

Due to limited computational resources, evaluation results are extracted from the original papers of several models if the same test set is used with similar setting, as denoted in the result tables. In stage1, we compared our models with ProtST and ProtT3. In stage2, we compare with Galactica, Prot2Text-V2, and ProtT3. Codes are not available for ProtChatGPT, and we do not have enough computational resources to re-train Prot2Chat on our dataset.

### E. Results of Stage 1: Protein–Text Retrieval

Table II shows stage1 evaluation results on the protein– text retrieval task after 50 epochs’ training. ProtBLIP2-SST improves test-set Acc and Rec@20 over ProtT3 at matched parameter scales. ProtBLIP2-SST (650M) achieves the highest average score. ProtT3 employs sequence-only Esm2 encoders without structural information. ProstT5 gains implicit structure awareness from pre-training and takes only sequence input. These factors limit their ability to capture fold-level functional constraints. These results demonstrate that integrating explicit structural information with sequence information effectively improves protein–text modeling.

**TABLE II.** Stage1 protein–text retrieval. *in-batch* metrics are computed on validation mini-batches; *test set* metrics use the alphafoldDB-filtered 9,239-protein test split with ptm re-ranking over 128 ptc candidates. Average is the mean of all eight Acc/Rec@20 numbers shown. * denote reported metrics extracted from the ProtT3 paper.

| Encoder | In-batch P→T |  | In-batch T→P |  | Test P→T |  | Test T→P |  | Average |
| --- | --- | --- | --- | --- | --- | --- | --- | --- | --- |
|  | Acc | R@20 | Acc | R@20 | Acc | R@20 | Acc | R@20 |  |
| *Baselines* |  |  |  |  |  |  |  |  |  |
| ProtST* | 81.40 | 99.40 | 79.60 | 99.40 | 15.90 | 56.60 | 17.80 | 63.50 | 64.20 |
| ProtT3 (35M) | 96.16 | 99.88 | 96.02 | 99.81 | 64.48 | 93.97 | 63.56 | 93.62 | 88.44 |
| ProtT3 (650M) | 97.00 | 99.86 | 97.01 | 99.88 | 67.68 | 95.14 | 67.49 | 95.27 | 89.92 |
| BLIP2 + ProstT5 (1.5B) | 97.37 | 99.94 | 97.14 | 99.91 | 66.75 | 96.21 | 66.99 | 95.89 | 90.03 |
| *ProtBLIP2-SST (35M) ablations* |  |  |  |  |  |  |  |  |  |
| w/o PTM | 96.46 | 99.47 | 96.13 | 99.36 | 68.75 | 95.18 | 64.95 | 94.48 | 89.35 |
| w/o PCap | 96.21 | 99.73 | 95.88 | 99.72 | 63.85 | 94.32 | 63.35 | 94.29 | 88.42 |
| Full model | 97.10 | 99.92 | 96.72 | 99.88 | 66.34 | 95.56 | 65.47 | 95.22 | 89.53 |
| *ProtBLIP2-SST (650M)* |  |  |  |  |  |  |  |  |  |
| Full model | **97.94** | **99.96** | **97.64** | **99.91** | **69.28** | **96.75** | **68.70** | **96.49** | **90.83** |

Due to limited computational resources, ablation studies are conducted exclusively on the 35M model. Removing PCap or PTM results in performance drop on average and removing PCap causes the largest degradation, confirming that the captioning objective is critical for extracting text-relevant protein features. Surprisingly, removing PTM improves test-set protein-to-text accuracy by 2.41%, suggesting room for more efficient and robust alignment objectives design.

### F. Results of Stage 2: Protein Caption Generation

Table III shows that ProtBLIP2-SST with integrated sequence and structure information achieves the best in BLEU, ROUGE, and METEOR, and second best in Exact match among existing models. ProtT3 (650M) is close but consistently lower. ProtBLIP2-SST 35M improves over ProtT3(35M) on BLEU-4 (+0.63), ROUGE-L (+0.97), and METEOR (+0.63) with comparable exact match (30.59 vs. 30.33).

**TABLE III.** Stage2 Swiss-Prot captioning on the test set. * denote reported metrics extracted from the original paper.

| Model | EM | B-2 | B-4 | R-1 | R-2 | R-L | Mtr |
| --- | --- | --- | --- | --- | --- | --- | --- |
| LoRA finetune (Galactica) | 13.49 | 42.48 | 37.79 | 44.42 | 34.73 | 42.46 | 48.27 |
| ProtT3 (35M) | 30.33 | 57.86 | 54.74 | 65.60 | 59.28 | 64.29 | 65.99 |
| ProtBLIP2-SST (35M) | 30.59 | 58.33 | 55.37 | 66.50 | 60.51 | 65.26 | 66.62 |
| ProtT3 (650M) | 32.83 | 61.01 | 58.12 | 68.94 | 63.15 | 67.77 | 69.10 |
| Prot2Text-V2* | **39.16** | 46.67 | 43.34 | 57.24 | 50.17 | 55.39 | 64.58 |
| ProtBLIP2-SST (650M) | 33.07 | **61.41** | **58.53** | **69.38** | **63.68** | **68.23** | **69.54** |

Table IV shows ablation test results on alignment and architectural choices at the 35M scale. We observe that Q-Former alignment (ProtT3/ProtBLIP2-SST) substantially outperforms feeding unaligned Esm2 hidden states from the last layer of all residues directly to Galactica, indicating alignment improves LLM fine-tuning. Simple LoRA fine-tuning of Galactica on raw amino-acid (AA) strings reaches only 24.4% exact match, and zero-shot baselines fail entirely. We notice that the scientific LLM Galactica uses special marker tokens (eg. [START AMINO] and [END AMINO]) to denote proteins [7]. However, improvement is not achieved by using these special markers (denoted as marker+AA).

**TABLE IV.** Stage2 ablation tests on ProtBlip2-SST (35M) on the test set.

| Variant | EM | B-2 | B-4 | R-1 | R-2 | R-L | Mtr |
| --- | --- | --- | --- | --- | --- | --- | --- |
| No finetune (marker + AA) | 0.00 | 1.94 | 0.58 | 7.33 | 1.89 | 6.20 | 5.11 |
| LoRA finetune (raw AA) | 24.43 | 40.72 | 37.15 | 54.67 | 46.49 | 53.05 | 57.40 |
| LoRA finetune (marker + AA) | 24.22 | 40.54 | 36.97 | 54.60 | 46.41 | 53.01 | 57.16 |
| Esm2 hidden states (no alignment) | 30.92 | 46.55 | 43.66 | 64.13 | 57.90 | 62.91 | 64.51 |
| ProtT3 (Esm2 + 8 queries) | 30.33 | 57.86 | 54.74 | 65.60 | 59.28 | 64.29 | 65.99 |
| ProtBLIP2-SST (SaProt + 8 queries) | **30.59** | **58.33** | **55.37** | **66.50** | **60.51** | **65.26** | **66.62** |

### G. Qualitative Analysis

Side-by-side analysis on all 9,239 test samples shows ProtBLIP2-SST wins on extact match cases than ProtT3 at both 35M (+24 wins: ProtBLIP2-SST achieves exact match but ProtT3 does not) and 650M (+22 wins) (Table V). In addition to exact match, we also calculate token-F1 scores as a less strict metric to facilitate field-stratified analysis. Token-F1 is a per-sample score that measures word overlap between a predicted caption and its ground truth based on their set of words. Table VI shows that ProtBLIP2-SST, which integrates structural information, achieves more token-F1 wins when predicting FUNCTION descriptions that rely on structural context, such as identifying specific pathways or enzyme targets. An example is shown in Figure 4. In contrast, ProtT3, which infers protein function from sequence alone, performs better on short *SIMILARITY* family labels driven by sequence motifs (e.g., HSP70, asp23). We found that both models inherit Galactica’s preference to produce predictions beginning with *FUNCTION* even when ground truth begins with *SIMILARITY*, suggesting future work on field-balanced decoding strategies.

**TABLE V.** Exact match comparison between ProtBlip2-SST and ProtT3.

| Scale | ProtBLIP2-SST<br>Exact only | ProtT3<br>Exact only | Both<br>Exact | Neither<br>Exact |
| --- | --- | --- | --- | --- |
| 35M | **316** | 292 | 2,510 | 6,121 |
| 650M | **283** | 261 | 2,772 | 5,923 |

**TABLE VI.**
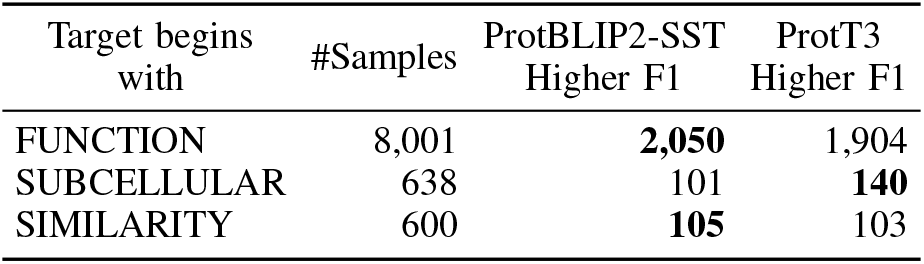
650M scale model comparison by description field.

**Fig. 4.**
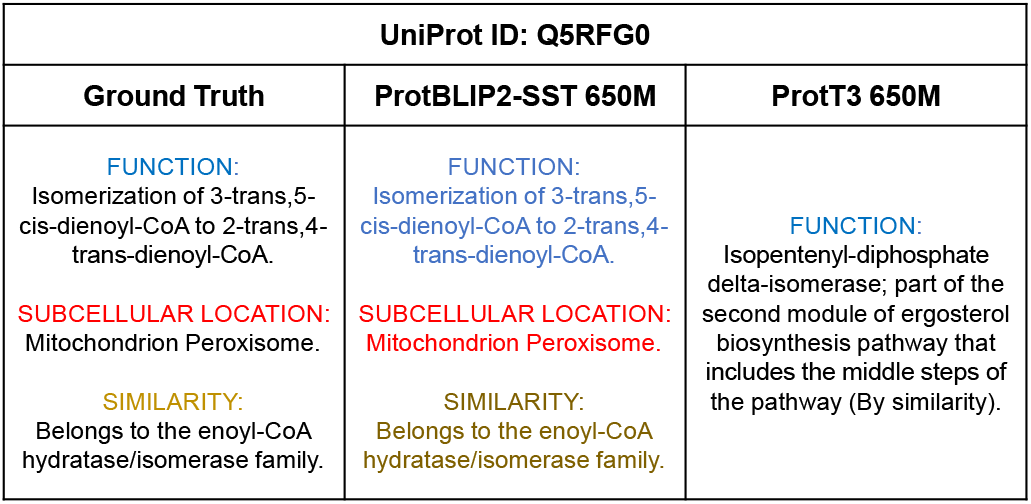
An example of ProtBLIP2-SST outperforming ProtT3 for captioning.

## V. Conclusion

In this study, we introduce ProtBLIP2-SST, a unified framework that integrates protein sequence, structure, and text for open-ended protein function captioning. Utilizing SaProt as a structure-aware protein encoder and Q-Former for cross-modal alignment, our model achieves consistent improvements over other sequence-only models on both retrieval and captioning tasks. Our field-stratified analysis reveals an important insight: the performance gains from incorporating 3D structural information are most significant for FUNCTION descriptions, where fold- and active-site-level constraints help distinguish functional mechanisms. In contrast, SIMILARITY labels are largely driven by sequence motifs. In addition, we observe that Galactica exhibits a strong preference for generating FUNCTION-oriented responses even when the ground-truth caption begins with SUBCELLULAR LOCATION or SIMILARITY. These observations suggest that gains from structural information are task-dependent and LLM-dependent. Future work includes designing more efficient alignment objectives and field-balanced decoding strategies for the LLM decoder.

## Notes

### Competing Interest Statement

The authors have declared no competing interest.

## References

[1] S. A. Aleksander, J. Balhoff, S. Carbon, J. M. Cherry, J. Drabkin, D. Ebert, M. Feuermann, P. Gaudet, N. L. Harris, et al., “The gene ontology knowledgebase in 2023,” Genetics, vol. 224, no. 1, p. iyad031, 2023.

[2] N. Zhou, Y. Jiang, T. R. Bergquist, A. J. Lee, B. Z. Kacsoh, A. W. Crocker, K. A. Lewis, G. Georghiou, H. N. Nguyen, M. N. Hamid, et al., “The cafa challenge reports improved protein function prediction and new functional annotations for hundreds of genes through experimental screens,” Genome Biol., vol. 20, pp. 1–23, 2019.

[3] Y.-H. Zhu, C. Zhang, D.-J. Yu, and Y. Zhang, “Integrating unsupervised language model with triplet neural networks for protein gene ontology prediction,” PLOS Computational Biology, vol. 18, no. 12, p. e1010793, 2022.

[4] G. B. Oliveira, H. Pedrini, and Z. Dias, “Temprot: protein function annotation using transformers embeddings and homology search,” BMC bioinformatics, vol. 24, no. 1, p. 242, 2023.

[5] T. U. Consortium, “Uniprot: the universal protein knowledgebase in 2023,” Nucleic Acids Res., vol. 51, no. D1, pp. D523–D531, 2023.

[6] Q. Zhang, K. Ding, T. Lv, X. Wang, Q. Yin, Y. Zhang, J. Yu, Y. Wang, X. Li, Z. Xiang, et al., “Scientific large language models: A survey on biological & chemical domains,” ACM Computing Surveys, vol. 57, no. 6, pp. 1–38, 2025.

[7] R. Taylor, M. Kardas, G. Cucurull, T. Scialom Hartshorn, E. Saravia, A. Poulton, V. Kerkez, and R. Stojnic, “Galactica: A large language model for science,” arXiv preprint arXiv:2211.09085, 2022.

[8] M. Xu, X. Yuan, S. Miret, and J. Tang, “ProtST: Multi-Modality Learning of Protein Sequences and Biomedical Texts,” in Proceedings of the 40th International Conference on Machine Learning.

[9] H. Zhou, M. Yin, W. Wu, M. Li, K. Fu, J. Chen, J. Wu, and Z. Wang, “Protclip: Function-informed protein multimodal learning,” in Proceedings of the AAAI Conference on Artificial Intelligence, vol. 39, pp. 22937–22945, 2025.

[10] A. Jararweh, O. Macaulay, D. Arredondo, Y. Hu, L. E. Tafoya, K. Virupakshappa, and A. Sahu, “Protein2text: Resampling mechanism to translate protein sequences into human-interpretable text,” in Proceedings of the 2025 Conference of the Nations of the Americas Chapter of the Association for Computational Linguistics: Human Language Technologies (Volume 3: Industry Track), pp. 918–937, 2025.

[11] Z. Liu, A. Zhang, H. Fei, E. Zhang, X. Wang, K. Kawaguchi, and T.-S. Chua, “Prott3: Protein-to-text generation for text-based protein understanding,” in Proceedings of the 62nd Annual Meeting of the Association for Computational Linguistics (Volume 1: Long Papers), pp. 5949–5966, 2024.

[12] M. Heinzinger, K. Weissenow, J. G. Sanchez, A. Henkel, M. Mirdita, M. Steinegger, and B. Rost, “Bilingual language model for protein sequence and structure,” vol. 6, no. 4, p. lqae150.

[13] J. Ouyang-Zhang, C. Gong, Y. Zhao, P. Krähenbühl, Klivans, and D. Diaz, “Distilling structural representations into protein sequence models,” in International Conference on Learning Representations, vol. 2025, pp. 93715–93727, 2025.

[14] J. Su, C. Han, Y. Zhou, J. Shan, X. Zhou, and F. Yuan, “Saprot: Protein language modeling with structure-aware vocabulary,” in International Conference on Learning Representations, vol. 2024, pp. 6987–7009, 2024.

[15] T. Hayes, R. Rao, H. Akin, N. J. Sofroniew, D. Oktay, Z. Lin, R. Verkuil, V. Q. Tran, J. Deaton, M. Wiggert, et al., “Simulating 500 million years of evolution with a language model,” Science, vol. 387, no. 6736, pp. 850–858, 2025.

[16] J. Su, Y. He, S. You, S. Jiang, X. Zhou, X. Zhang, Y. Wang, X. Su, I. Tolstoy, X. Chang, H. Lu, and F. Yuan, “A trimodal protein language model enables advanced protein searches,” pp. 1–7.

[17] H. Guo, M. Huo, R. Zhang, and P. Xie, “Proteinchat: Towards achieving chatgpt-like functionalities on protein 3d structures,” Authorea Preprints, 2023.

[18] Z. Wang, Z. Ma, Z. Cao, C. Zhou, J. Zhang, and Y. Gao, “Prot2Chat: Protein LLM with Early-Fusion of Text, Sequence and Structure.”

[19] C. Wang, H. Fan, R. Quan, L. Yao, and Y. Yang, “ProtChatGPT: Towards Understanding Proteins with Hybrid Representation and Large Language Models,” in Proceedings of the 48th International ACM SIGIR Conference on Research and Development in Information Retrieval, SIGIR ‘25, pp. 1076–1086, Association for Computing Machinery.

[20] J. Li, D. Li, S. Savarese, and S. Hoi, “BLIP-2: Bootstrapping Language-Image Pre-training with Frozen Image Encoders and Large Language Models,” in Proceedings of the 40th International Conference on Machine Learning.

[21] M. Varadi, D. Bertoni, P. Magana, U. Paramval Pidruchna, M. Radhakrishnan, M. Tsenkov, S. Nair, M. Mirdita, J. Yeo, et al., “Alphafold protein structure database in 2024: providing structure coverage for over 214 million protein sequences,” Nucleic Acids Res., vol. 52, no. D1, pp. D368–D375, 2024.

[22] M. Van Kempen, S. S. Kim, C. Tumescheit, M. Mirdita, Lee, C. L. Gilchrist, J. Söding, and M. Steinegger, “Fast and accurate protein structure search with fold-seek,” Nat. Biotechnol., vol. 42, no. 2, pp. 243–246, 2024.

[23] Z. Lin, H. Akin, R. Rao, B. Hie, Z. Zhu, W. Lu, A. dos Santos Costa, M. Fazel-Zarandi, T. Sercu, S. Candido, et al., “Language models of protein sequences at the scale of evolution enable accurate structure prediction,” BioRxiv, vol. 2022, p. 500902, 2022.

[24] A. Radford, J. W. Kim, C. Hallacy, A. Ramesh, G. Goh, S. Agarwal, G. Sastry, A. Askell, P. Mishkin, J. Clark, et al., “Learning transferable visual models from natural language supervision,” in International conference on machine learning, pp. 8748–8763, PmLR, 2021.

[25] J. Li, D. Li, C. Xiong, and S. Hoi, “Blip: Bootstrapping language-image pre-training for unified vision-language understanding and generation,” in International conference on machine learning, pp. 12888–12900, PMLR, 2022.

[26] E. J. Hu, Y. Shen, P. Wallis, Z. Allen-Zhu, Y. Li, S. Wang, Wang, W. Chen, et al., “Lora: Low-rank adaptation of large language models.,” Iclr, vol. 1, no. 2, p. 3, 2022.

[27] J. Dauparas, I. Anishchenko, N. Bennett, H. Bai, R. J. Ragotte, L. F. Milles, B. I. Wicky, A. Courbet, R. J. de Haas, N. Bethel, et al., “Robust deep learning–based protein sequence design using proteinmpnn,” Science, vol. 378, no. 6615, pp. 49–56, 2022.

[28] Y. Gu, R. Tinn, H. Cheng, M. Lucas, N. Usuyama, X. Liu, T. Naumann, J. Gao, and H. Poon, “Domain-specific language model pretraining for biomedical natural language processing,” ACM Transactions on Computing for Healthcare (HEALTH), vol. 3, no. 1, pp. 1–23, 2021.

